# EIF4A3 associated splicing and nonsense mediated decay defined by a systems analysis with novel small molecule inhibitors

**DOI:** 10.1101/189639

**Authors:** Alborz Mazloomian, Shinsuke Araki, Momoko Ohori, Damian Yap, Shoichi Nakao, Atsushi Nakanishi, Sohrab Shah, Samuel Aparicio

## Abstract

Chemical biology approaches to the global functions of splicing reactions are gaining momentum, with an increasing repertoire of small molecule probes becoming available. Here we map the association of eIF4A3 with transcript expression, NMD and alternative splicing using a set of selective novel small molecule allosteric helicase inhibitors whose synthesis and chemical properties we have recently described. We show through analysis of dose monotonic transcriptional responses to increasing inhibition that both full length and NMD prone transcripts link eIF4A3 to normal functioning of cell division including chromosome segregation and cell cycle checkpoints, pointing to a conserved role of splicing and transcript quality processing in cell cycle functions. Cell cycle analysis and microscopy of inhibitor treated cells demonstrates chromosome mis-segregation and spindle defects, associated with a G2/M arrest, validating this observation. Through analysis of conserved alternative splicing patterns exhibiting monotonic responses, we find that eIF4A3 dependent alternative splicing involves exons that are longer and introns that are shorter than transcripts not modulated by eIF4A3. Moreover we observe conservation of over/under representation of RBP binding motif density over introns and exons implicated eIF4A3 modulated skipped exon and retained introns. The distribution of motif densities over 5’ and branch intron sites and 5’ exons is consistent with function of the exon-junction complex. Taken together we have defined a fraction of the transcriptome dependent on eIF4A3 functions and revealed a link between eIF4A3 and cell cycle regulation. The systems approach described here suggests additional avenues for therapeutic exploitation of eIF4A3 functions in cancer and related diseases.

## Introduction

Chemical biology approaches to the global functions of splicing reactions are gaining momentum, with an increasing repertoire of small molecule probes becoming available. Here we define the global transriptomic functions of eIF4A3 using a set of selective novel small molecule allosteric helicase inhibitors whose synthesis and chemical properties we have recently described (Ito *et al*, 2017b,a)).

The eukaryotic initiation factor 4A (eIF4A) belongs to the Asp-Glu-Ala-Asp (DEAD) box superfamily (SF) of proteins that have ATP-dependent RNA helicase activity and are involved in various aspects of RNA biology, from transcription and translation to mRNA decay (Linder and Jankowsky, 2011). There are three paralogous genes of eIF4A, eIF4A1,2,3. Although eIF4A3 exhibits strong phylogenetic conservation between species and high homology at the amino-acid level to the translation initiation factors, eIF4A1 and eIF4A2, the confusingly-named eIF4A3 (also known as DDX48, Nuk34, and hNMP 265) is actually a core component of the exon junction complex (EJC) (Li *et al*, 1999; Chan *et al*, 2004), the assembly of which is closely associated with splicing and does not play a significant role in translation initiation, in contrast to eIF4A1 and eIF4A2. The splicing factor CWC22 is essential for the initial formation of the EJC through the recruitment of eIF4A3 to spliceosomes (Barbosa *et al*, 2012), attesting to the close association between splicing and EJC formation. The binding of MAGOH and Y14 to eIF4A3 locks the pre-EJC onto the mRNA (Nielsen *et al*, 2009) while the incorporation of MLN51 (for Metastatic Lymph Node 51, also known as hBarentsz (BTZ) or CASC3) (Buchwald *et al*, 2010) completes the core EJC (Le Hir and Andersen, 2008). Other members of the multiprotein EJC, include SRm160/SRRM1, DEK, RNPS1 and ALY/REF (Le Hir *et al*, 2000). The EJC is deposited by the spliceosome on the 5' exon, 20-24 nucleotides (nt) upstream of the recently spliced out intron, hence marking the location of exon-exon boundaries on mRNA (Le Hir *et al*, 2000). This is significant given the previous evidence that nonsense codons (generating premature stops hence called premature stop codons, PTCs) located in the last exon rarely elicit nonsense-mediated mRNA decay (NMD) (Hall and Thein, 1994). Furthermore, NMD is triggered if the PTC is at least 50-55 nt upstream of the final exon-exon junction (Hentze and Kulozik, 1999). Therefore, memory of the exon junctions is crucial to the NMD process and the EJC is suitably placed to confer such a function. Indeed, EJC complex members, eIF4A3 (Wang *et al*, 2014), RNPS1 (Lykke-Andersen *et al*, 2001), UPF1-3 (Lykke-Andersen *et al*, 2000), Y14 (Gehring *et al*, 2003), MLN51 (Wang *et al*, 2014) have been shown to play key roles in NMD and independent deletion of eIF4A3, Y14, MLN51, respectively, has been shown to affect an overlapping set of genes that are potentially regulated by NMD (Wang *et al*, 2014). Previous work using whole genome approaches have shown that block of NMD through specific knock down of UPF1 in yeast leads to the modulation of expression of NMD transcripts (Mendell *et al*, 2004).

NMD appears to be an evolutionarily conserved surveillance mechanism monitoring eukaryotic mRNA translation and targeting mRNAs with PTCs for rapid degradation thereby averting the energetically wasteful or potentially even deleterious effects of the accumulation of truncated polypeptides. It has been recently recognized that NMD also plays a fundamental role in the physiological regulation of gene expression from wild type mRNA (reviewed in (He and Jacobson, 2015)). Well-studied examples include the splicing-dependent inclusion of PTC-containing exons or 3’UTRs in the transcripts of SR family of factors that are then regulated by NMD (Lareau *et al*, 2007). NMD function is also involved in diverse cellular processes, from cell growth and proliferation, to development and differentiation, from innate immunity, antiviral or stress responses, to neuronal activity or behaviour (reviewed in (He and Jacobson, 2015)).

Mutations in eIF4A3 have been found in Richieri-Costa-Pereira syndrome (RCPS), an autosomal-recessive acrofacial dysostosis characterized by craniofacial anomalies and severe limb defects (Favaro *et al*, 2014). These patients had an expanded number of the repeat motifs in the 5’UTR of eIF4A3 as compared with control unaffected individuals. These affected the levels of the eIF4A3 transcript but apparently not its splicing. An unrelated patient with an atypical presentation of RCPS, had a normal 5’UTR but a missense mutation in the exonic region which was proposed to affect the interaction of eIF4A3 to UPF3B. Furthermore, zebrafish embryos with depleted levels of eIF4a3 also exhibited a craniofacial phenotype.

Here we show that graded-pharmacological specific inhibition of eIF4A3, a core component of the EJC, results in class-specific splicing defects and the monotonic increase of NMD prone transcripts of genes which are involved in a range of specific functions, including cell cycle (particularly G2/M checkpoint) as well as chromosomal alignment. An unexpected finding is the fact that not all alternative spliced transcripts generated by eIF4A3 (and perhaps EJC) inhibition resulted in the generation of NMD prone isoforms and vice versa, indicating distinct roles for eIF4A3 and hence the EJC complex in splicing and NMD. To demonstrate the potential utility of the small molecule inhibitor of eIF4A3, we show that treatment of cells with the drug, results in spindle defects at the G2/M checkpoint as well as multiple chromosomal segregation defects. We further characterize the motifs that might suggest how eIF4A3 might act to accomplish these effects.

## Results

### Defining core eIF4A3 dependent transcriptional responses and NMD-prone transcripts with chemical probes

We (Funnell *et al*, 2017a) and others have used graded short duration pharmacological modulation of gene function with novel small molecule inhibitors to identify transcripts and alternative isoforms influenced by spliceosome components. Here we set out to define the transcriptional and splicing dependencies of the eIF4A3 helicase, a key component of the exon junction complex (EJC) using graded pharmacological inhibition with our recently discovered eIF4A3 small molecule inhibitors. For this purpose we contrasted two specific and active allosteric eIF4A3 inhibitors (T-595, T-202) that have similar scaffolds, with an inactive (>120 fold less active) but chemically identical steroisomer compound (T-598) as a control. We have recently described the structures and synthesis of these compounds: T-595 ((3S)-4-(4-Bromobenzoyl)-3-(4-chlorophenyl)piperazin-1-yl)(6-bromopyrazolo[1,5-a]pyridin-3-yl)methanone], compound 52a in (Ito *et al*, 2017b) T-595, compound 52b T-598) and T-202 a related scaffold (3-(4-(((3S)-4-(4-Bromobenzoyl)-3-(4-chlorophenyl)piperazin-1-yl)carbonyl)-5-methyl-1H-pyrazol-1-yl)benzonitrile, compound 53a)(Ito *et al*, 2017b). Both T-595 and T-202 are potent and specific (selectivity over EIF4A1,2; Supplementary Data 1 and (Ito *et al*, 2017b,a)) allosteric eutomer inhibitors of eIF4A3 activity in helicase unwinding assays and have been demonstrated to suppress nonsense mediated decay (NMD) in reporter assays (Iwatani-Yoshihara *et al*, 2017), however the global transcriptional and splicing consequences of eIF4A3 pharmacological inhibition have not been described.

We collected RNA-seq reads from cells treated with (Materials and Methods, Supplementary Table 1, Supplementary Figure 1) increasing concentrations of short duration exposure (6 hours, as in (Funnell *et al*, 2017a)) to each of the three compounds, from two mammalian cell lines HCT116 and HeLa, generating 32 sequencing libraries. RNA-seq reads were aligned and quantified as previously described (Materials and Methods; Supplementary Figure 1). We first examined eIF4A3 trancriptional responses, employing weighted correlation network analysis (WGCNA) (Langfelder and Horvath, 2008) across control and increasing doses of active and control compounds, for normalized RNA expression (Materials and Methods) as we have demonstrated for other splicing factors (Funnell *et al*, 2017a). Consistent with the chemical and pharmacological potency of the compounds, treatment of cells with both active eutomer allosteric inhibitors T-595 and T-202 resulted in a large number of monotonically increasing (T-595 n=1244, T-202 n=1417) or decreasing (T-595 n=1885, T-202 n=1341) transcripts (Figure 1a) in response to increasing inhibition, whereas treatment with the inactive distomer T-598 gave much smaller clusters of transcripts that showed no clear monotonic dose relationship. A high dynamic range of FPKM values was observed for monotonically increasing gene transcripts (0-5907), and monotonically decreasing genes (0-7958), across all compound:cell line pairs (Supplementary Figure 2).

**Figure 1:**
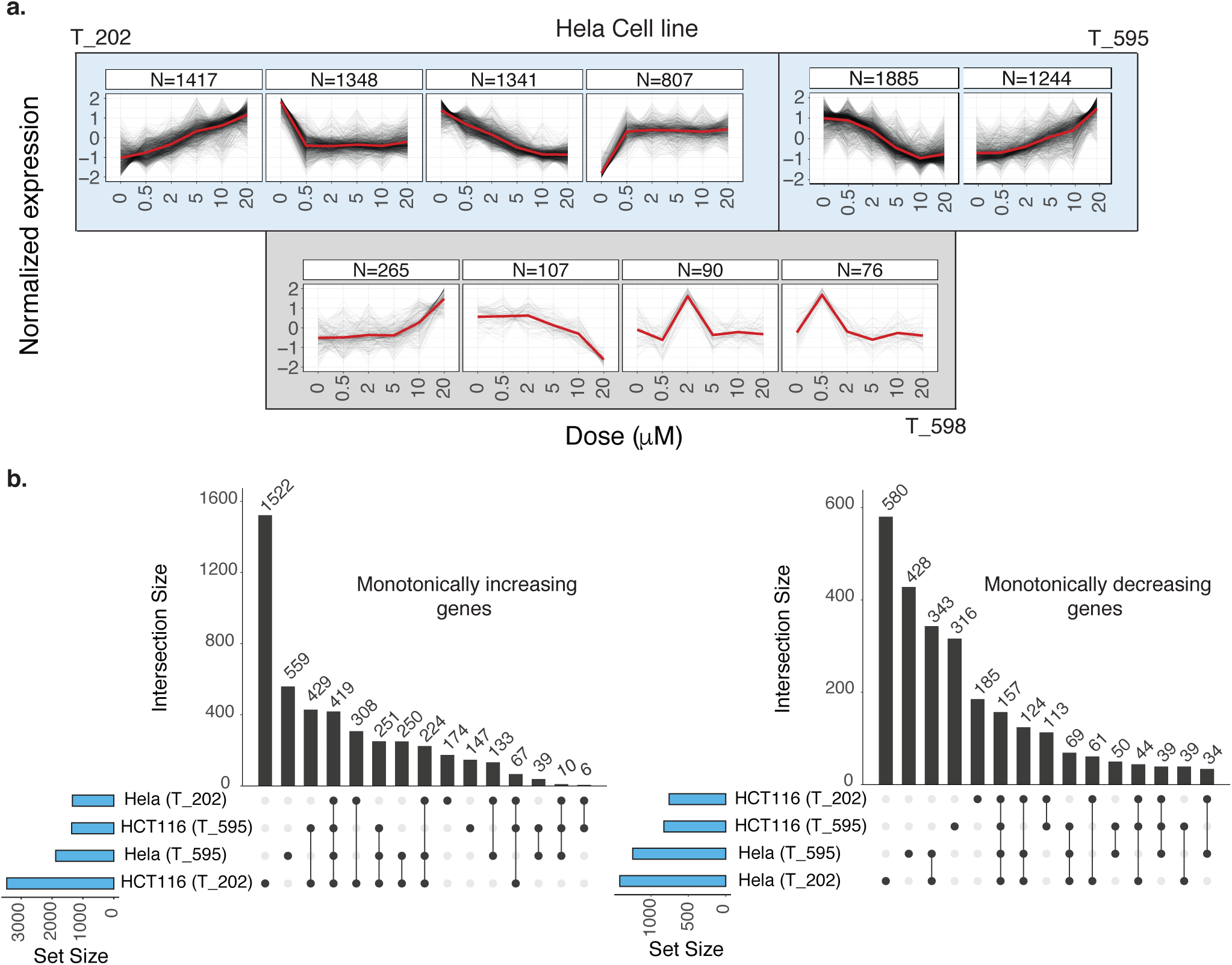
Determining core transcriptional response genes. **a.** Results of clustering gene expression responses using WGCNA. Blue and gray background colors represent the clustering of gene expression profiles for the active compounds and the control compound, respectively (at most 4 clusters are shown per each compound, sorted based on cluster sizes). X and Y axes represent inhibitor concentrations and gene expression values normalized using upper quartile normalization method. Each black line illustrates the expression response of one gene in a given condition, and the red lines represent the consensus response in each cluster computed by averaging expression values of all the genes. **b.** Overlap sizes between subsets of monotonically increasing/decreasing responses are shown for the two active compounds (T-202 and T-595). Each bar corresponds to the set of events only present in the cell line:inhibitor pairs determined by black circles.

Across two cell lines and the two active eutomers, monotonically increasing transcripts (n=4538) dominate over monotonically decreasing (n=2582) (Figure 1b, Supplementary Data 2, 3) and in both cases a large proportion (∼42% and ∼47%) of monotonic responses were shared in common between two or more conditions across cell lines and compounds, indicating a high degree of conservation in eIF4A3 dependent transcriptional responses.

The role of eIF4A3 in exon-junction complex activity and its involvement in NMD prompted us to ask whether we could measure a global increase in NMD prone transcripts when eIF4A3 is pharmacologically inhibited. To address this, we utilised Ensembl annotations (Cunningham *et al*, 2015) for NMD prone transcript isoforms and re-examined the inhibition-response relationship using WGCNA. A large number of monotonically increasing NMD prone transcripts were found on treatments with both the active compounds (T-595 n=1168, T-202 n=1347), whereas no clear monotonically increasing cluster was observed with treatments with the chemically identical T-598 isomer (Figure 2a, Supplementary Data 4). Moreover, no large monotonically decreasingly clusters were observed in any of the conditions, when only NMD prone isoforms were considered, entirely consistent fact that under normal treatment without eIF4A3 inhibiting drugs, NMD-prone transcripts are degraded. Although isoform ratios are harder to measure accurately, we also examined the ratio of NMD expression to corresponding genes (Figure 2b) and similarly observed the largest dominant clusters for both compounds represented monotonically increases of NMD prone isoforms over the genes, although clusters were overall smaller (T-595 n=620, T-202 n=535; Supplementary Data 5). This suggested that the specific NMD isoforms increase due to the block of NMD rather than the overall upregulation all exons of the gene. In contrast with the unstratified analysis of monotonic eIF4A3 response clusters above, the majority of monotonically responsive transcripts overlapped in 2 or more conditions of absolute (Figure 2c, n=1437/2240, 64%) and isoform ratio comparisons also overlapped to a large degree (Figure 2d, n=609/1368, 45%). These data are consistent with the dose-dependent inhibition of eIF4A3 by the eutomers resulting in a conserved (across different cell lines) monotonic increase in NMD prone transcripts.

**Figure 2:**
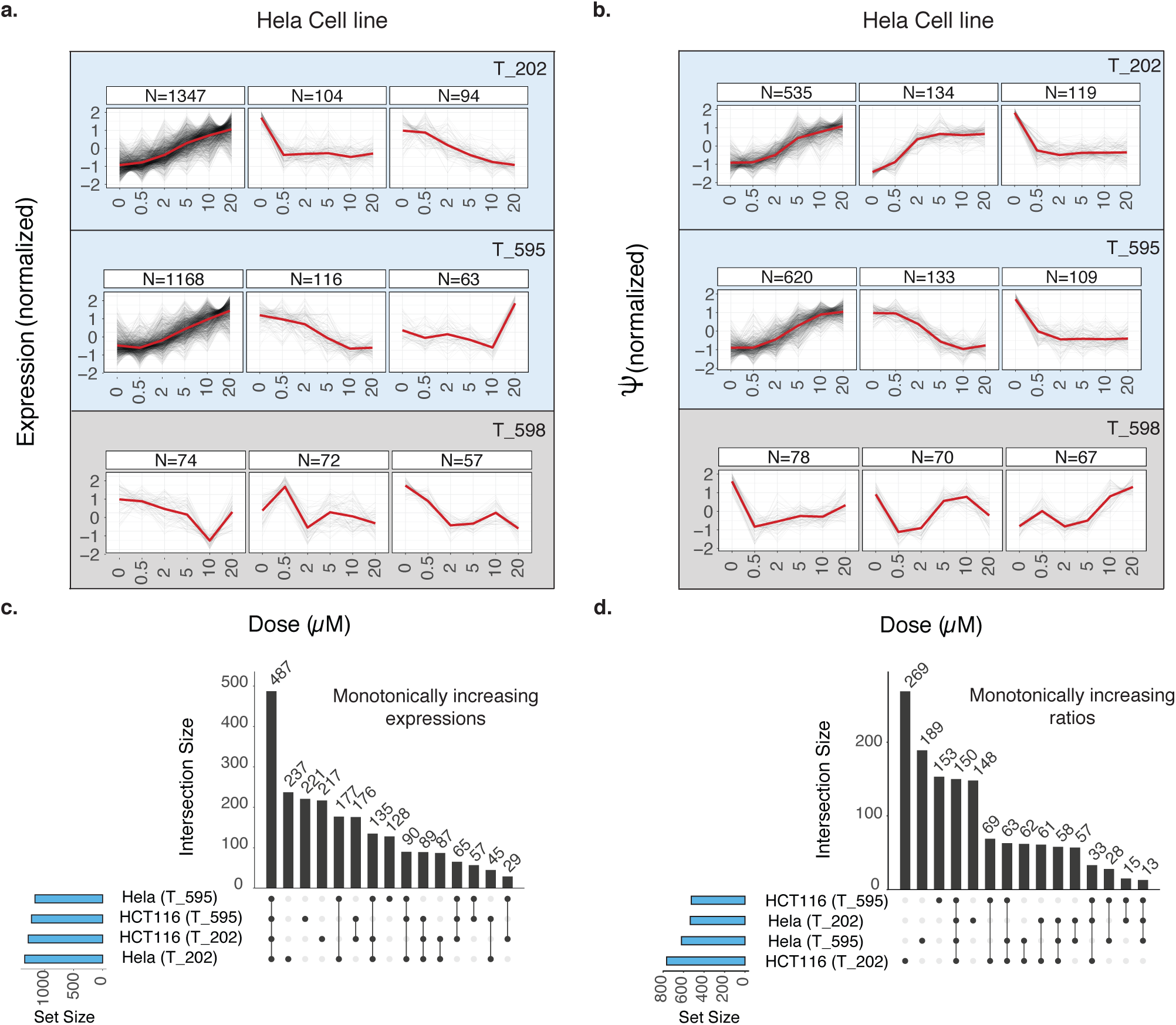
Investigating NMD prone transcripts responses to gradual eIF4A3 inhibition. **a.** Results of clustering expression profiles for NMD prone transcripts as annotated by Ensembl. Blue and gray background colors represent the clustering of isoform expression profiles for the active compounds and the control compound, respectively (3 top clusters are shown per each compound, sorted based on cluster sizes). X and Y axes represent inhibitor concentrations, and isoform expression values normalized using upper quartile normalization method. Each black line illustrates expression response of one NMD prone isoform in a given condition, and the red lines represent the consensus response in each cluster computed by averaging expression values of all the genes. Most isoforms are over-expressed in higher inhibition levels of active compounds, in contrast to the observed responses when cells are treated by the control compound. **b.** Inclusion levels of NMD prone isoforms (Ψ values normalized by the the expression of corresponding genes) are clustered using WGCNA. Similar to part a, isoforms are predominantly over-expressed for active compounds shown in blue backgrounds. **c., d.** Overlap sizes among subsets of monotonically increasing profiles of **a** and **b** are shown for the two active compounds. Each bar corresponds to the set of events only detected in the cell line:inhibitor pairs determined by black circles.

Taking the global transcript and NMD isoform analysis together, we noticed monotonically reduced transcription of the most abundant isoforms and/or increased NMD prone isoforms are evident among genes responsible for RNA processing/splicing (e.g. SRSF6, RBMX, SART3, SRSF7, SRSF3, SRSF2, HNRNPA1, RBM38, SFPQ, QKI, HNRNPL, MBNL2, PABPC4, RBM4B, PSPC1, RBM41, ESRP2) and cell cycle regulators/cell cycle checkpoints (e.g. CDK7, SMC1A, CDC27, CCNE2, CCNL2) (Supplementary Figures 3a, 4a). We therefore systematically investigated which biological processes (BP) were affected, through Gene Ontology (GO)-term pathway enrichment statistical analysis (EnrichmentMap (Merico *et al*, 2010a), Materials and Methods) of dose monotonic gene clusters (Figure 3a, Supplementary Data 6, 7, 8, 9). We observed a high degree of concordance between the two active compounds (Figure 3b) and also between the two cell lines, indicating that the overall correlation in eIF4A3 dependent cellular functions between cell types was relatively high, for both T-202 (s = 0.97, *r*^2^ = 0.65) and T-595 (s = 1.1, *r*^2^ = 0.87). Interestingly, both up and down regulated cell cycle checkpoint processes were observed, alongside down-regulated clusters enriched (FDR <0.01, hypergeometric test) for functions in cell cycle, cytokinesis, cell division, chromosome localization, spindle assembly/organization, DNA repair and protein localisation. These are all functions which can be associated with disordered regulation of cell division checkpoints. In contrast, up-regulated clusters were enriched (FDR <0.01, hypergeometric test) for stress responses, ER function, apoptosis components and chromatin modification functions.

**Figure 3:**
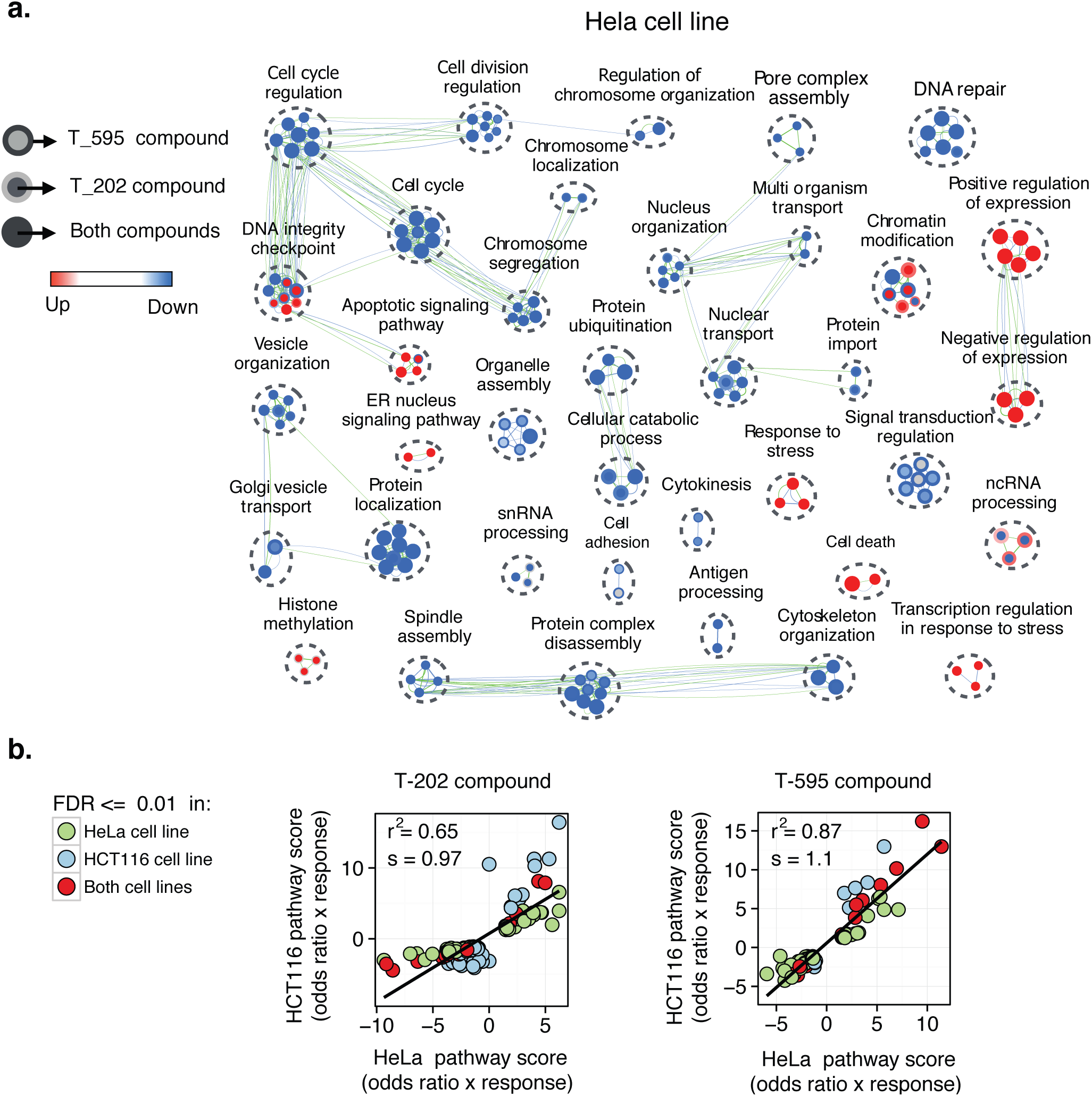
Biological processes affected most by eIF4A3 depletion. **a.** Enrichment map from GO term enrichment analysis of biological processes. Each node represents a biological process with genes over-represented in the set of monotonically increasing (up, red)/decreasing (down, blue) clusters for HeLa cell line. Node cores and node rings illustrate the results for T-202 and T-595 compounds, and green and blue edges indicate the overlap between the identified genes of gene sets for T-202 and T-595 compounds, respectively. Biological processes are clustered using EnrichmentMap (Merico *et al*, 2010a) based on the overlap of their monotonic genes. **b.** A comparison between the enrichment scores (x and y axes) of GO terms identified in T-202 and T-595 compound libraries with FDR values <0.01. Each point represents a BP set enriched in HeLa or HCT116 cell line data. Odds ratios are calculated by dividing the percentage of the genes in a given GO set that are identified by the percentage of all the genes that are identified. To compute the final scores, odd ratio scores are multiplied by ‐1 only if the enriched BP corresponds to the set of decreasing response patterns.

**Figure 4:**
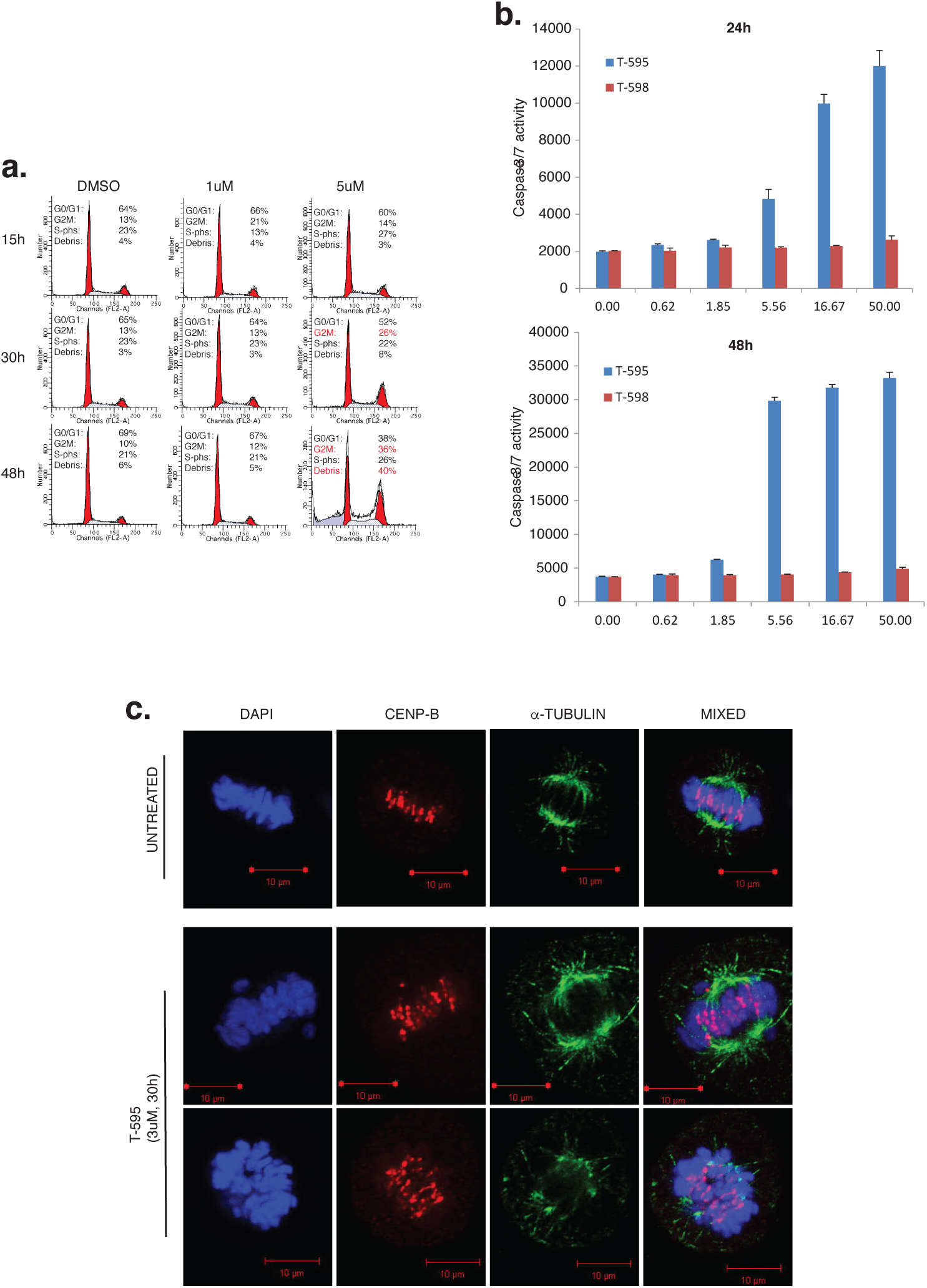
Biological effects of treatment with eIF4A3 inhibitor, T-595. **a.** Treatment with T-595 at high doses and after 48 hr results in G2M arrest and sub-G1/G0 fraction indicative of apoptosis. Cell cycle profiles as assessed by flow cytometry after the treatment with the respective compounds (DMSO, 1 or 5 *μ*M T-595) for the respective times shown. The percentage of cells assigned to be in the respective stages of the cell cycle and indicated on the respective charts. **b.** Apoptosis verified by treatment with eutomer, T-595 but not distomer, T-598. Graphs show independently measured apoptosis as assessed by Caspase 3/7 activity (vertical axis) when HeLa cells were treated for 24 hr with the respective concentrations of T-595 (blue bar) and T-598 (red bar) as indicated on the horizontal axis. **c.** Microscopy for spindle and chromosome proteins reveals chromosome mis-segregation. Representative images of untreated (top panel) and treated (5 *μ*M T-595 for 30 hr, bottom two panels) showing chromosomal DNA stained with DAPI (blue), centromere proteins stained with anti-CENP-B (red) or microtubules identified by anti-α-tubulin (green).

Similarly, we performed GO-term pathway enrichment analysis for the genes with NMD-prone transcripts that showed monotonic responses (Supplementary Figures 3a, 4a, Supplementary Data 10, 11, 12, 13, 14, 15, 16, 17). The isoform expression based analysis uncovered a larger number of biological process terms enriched by the set of genes with NMD-prone isoforms showing monotonic response patterns compared to isoform ratio based analysis. The enriched GO terms include cell cycle, cellular respiration, NMD, and protein localisation and some other pathways, some of which were also detected in the isoform ratio based analysis (DNA repair, RNA splicing and processing). There is again a notable concordance between the results of the two cell lines (Supplementary Figures 3b and 4b).

The systems analysis above strongly implicated cell cycle, spindle assembly, chromosome segregation, checkpoint processes that might be affected by reduced eIF4A3 function. To confirm in vitro, the biological relevance of the GO-term network analysis we measured the cell cycle responses and apoptotic fraction in response to increasing doses of the active and control compounds in both synchronised and unsynchronized cells (Figure 4a,b) by flow cytometry. Consistent with the network analysis, this revealed that prolonged and increasing inhibition of eIF4A3 (doses >5 *μ*M and >15 hours exposure) results in increased apoptosis, associated with a cell cycle arrest at the G2M boundary. Timed microscopy of cells (Figure 4c) with immunofluorescence staining for spindle and chromosome centrosome proteins (alpha-tubulin, CENP-B) revealed frequent chromosome mis-segregation and abnormal spindle assembly. Taken together, these data suggest that disrupted cell cycle associated with chromosome segregation abnormalities and a G2/M checkpoint induction, in addition to apoptosis are consequences of eIF4A3 inhibition.

### Defining eIF4A3 dependent alternative splicing (AS) events

The known functions of eIF4A3 in the EJC implicate both NMD and alternative splicing (AS) as regulated processes. To investigate eIF4A3 dependent AS regulation, we explored the graded inhibition RNA-seq data, with two computational methods that have partially overlapping but distinct AS feature sets (MISO and VAST-TOOLS), in order to obtain wide coverage of alternative splicing events. The mixture of isoforms (MISO) (Katz *et al*, 2010) framework applies a Bayesian approach to identify 8 AS types: skipped exons (SE), retained introns (RI), alternative first/last exons (AFE/ALE), alternative 3’/5’ splice sites (A3SS/A5SS), mutually exclusive exons (MXE), and tandem untranslated regions (TandemUTR). We separately examined each inhibitor:cell line pair (details in Materials and Methods).

The two active compounds induce dose-dependent increase in the total number of MISO determined AS events (Figure 5a). This trend is in contrast with the control compound, in T-598 treated cells, only a small total number of events were observed (161 events predicted more than once, as compared to 1405 and 788 events in T-202 and T-595 treated cells), and no dose-dependent trend could be observed. The average increase rate in the number of events between any two consecutive inhibitions for the active compounds (averaged over the two cell lines and the two compounds) ranges between 0.17 and 1.43, with the maximum increase at 5 *μ*M (compared to 2 *μ*M).

**Figure 5:**
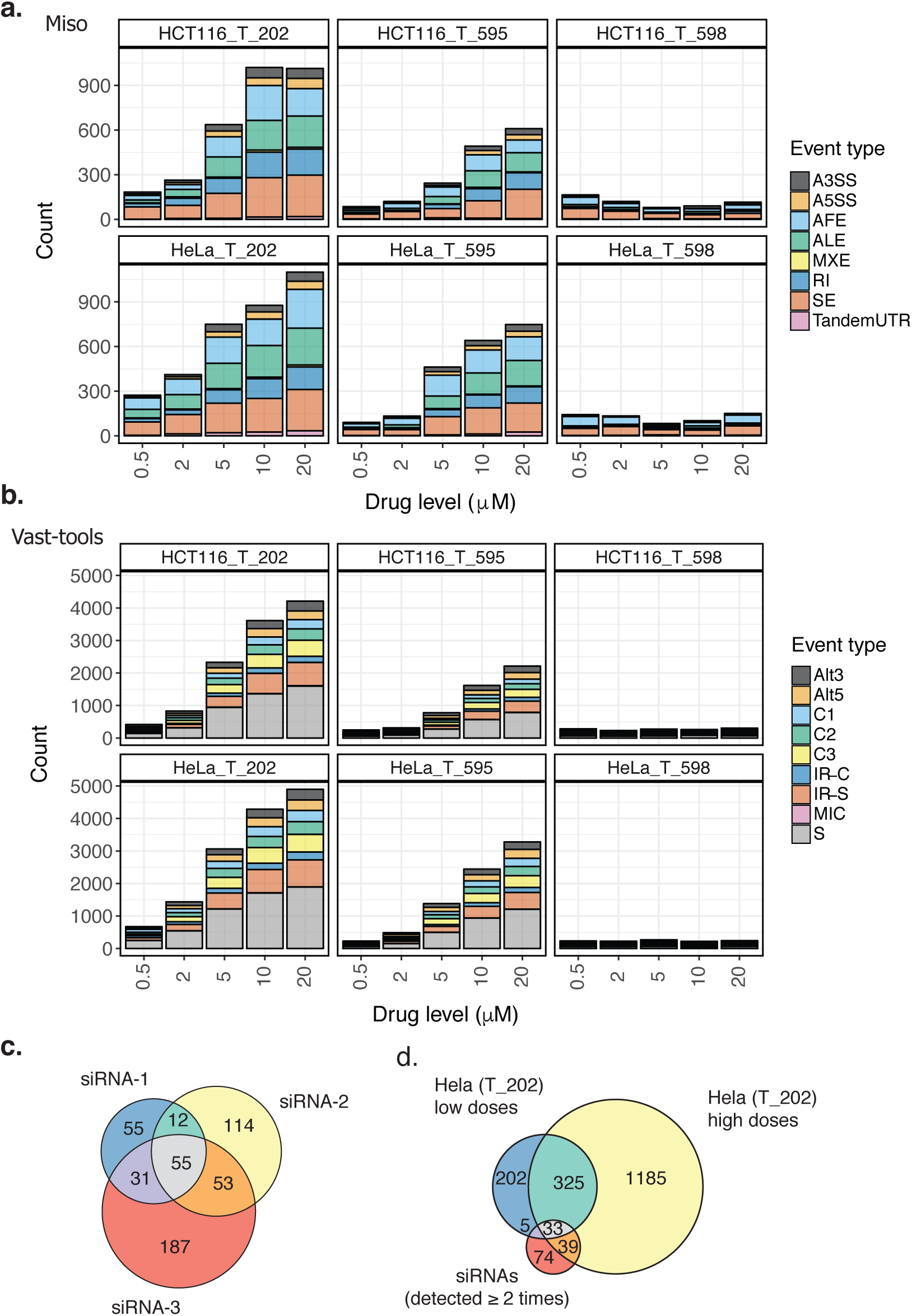
Characterizing AS events modulated by eIF4A3 inhibition. **a.** Counts of differentially spliced events (vertical axis) by MISO framework at each inhibitor concentration (horizontal axis) for each cell line:inhibitor condition. Only when cells were treated with active compounds, the number of events increased at higher drug concentrations. **b.** Similar patterns of increase in the number of identified AS events were observed when we used VAST-TOOLS. **c.** A venn diagram showing the overlap of MISO identified AS events among knock down experiments of eIF4A3 using 3 different siRNAs. **d.** A venn diagram displaying the overlap of MISO identified AS events when knocking down eIF4A3 with siRNAs (only events identified in at least two treatments are considered), treating the cells with low concentrations of T-202 inhibitor, and treating the cells with high concentrations of T-202 inhibitor.

SE is the most prevalent type of MISO observed AS event across multiple doses in both cell lines and the active inhibitors (N=1279 unique events). AFE (n=807), ALE (n=649), and RI (n=443) are the other abundant types of AS detected in our data sets. It should be noted however that only AS events previously annotated in MISO knowledge base are visible and the relative abundance of AS types are therefore likely influenced by how comprehensively they are annotated in the MISO knowledge base, and more generally, in their overall prevalence. Accordingly, we normalized the number of events detected in at least one condition for each event type by the total number of events of the same type in MISO knowledge base, and found the highest ratio for RI events (7.4% of all RI events, where the ratio ranges between 1.4% to 5.9% for the other AS types, Supplementary Table 2).

To extend our analysis, we also determined AS events using VAST-TOOLS (Irimia *et al*, 2014) (Figure 5b), which has additional feature types and a different event database underlying the method. VAST-TOOLS recognizes skipped exons, micro-exons, alternative 3’/5’ splice sites and is more sensitive to retained intron detection, in part due to the much larger annotation database (~155,000 vs ~6,000) of introns. Skipped exons are stratified into multiple groups based on the complexity of events (C1, C2, C3, S), and an alternative pipeline decides whether they should be classified as micro exons (MIC).

Similar to the MISO analysis, AS event counts increase with increasing concentrations of the active compounds, and most of the detected events are skipped exons (7040 unique events in all of libraries, Supplementary Table 3). More importantly the much larger annotation database (Supplementary Figure 5) of VAST-TOOLS enabled us to identify many more retained introns compared with MISO (2680 unique RI events). Also, more A3SS and A5SS events were reported by VAST-TOOLS compared to MISO (939 and 930 vs 250 and 185, respectively). A majority ~55% of MISO determined events overlapped with VAST-TOOLS determined events (Supplementary Figure 6).

We independently validated the eIF4A3 inhibition AS event trends by comparison with eIF4A3 siRNA knockdown experiments (Figure 5c, d). We treated HeLa cells with three siRNAs (siRNA-1, siRNA-2, and siRNA-3) directed to eIF4A3 transcripts, and one control siRNA (Supplementary Table 1). Treatment with siRNAs reduced eIF4A3 transcripts abundance by ~92% on average (Supplementary Figure 7). Approximately 31% of siRNA associated AS events were shared among at least two siRNAs. To compare these events with inhibitor associated events, we first grouped AS events in T-202 drug inhibition data into “low dose” (0.5 and 2 *μ*M) and “high dose”, based on the drug dose at which events were predicted. A large proportion of AS events in the low dose group were also found in the high dose group (63%), and 51% of the events predicted in more than two or more treatment:control siRNA comparison were also detected by drug inhibition. Moreover, the rate of overlap increases between low dose and high dose conditions. Similar patterns were observed when we compared siRNA results to T-595 compound (Supplementary Figure 8).

We clustered Ψ profiles of all the identified MISO AS events to characterize AS events showing responses in agreement with eIF4A3 inhibition levels (Figure 6a). The two dominant clusters in both cell lines for the two active compounds are sets of events with monotonically increasing (T-202 n=496, T-595 n=346) or monotonically decreasing (T-202 n=472, T-595 n=444) Ψ values as opposed to the control compound where no monotonic response is observed in clusters, and the cluster set sizes are much smaller. Similar trends were observed for HCT116 cell line libraries (Supplementary Figure 9). To determine the degree of conservation of eIF4A3 AS events we computed the overlap of monotonically increasing/decreasing events between the 4 drug:cell line conditions. Only a small fraction (Figure 6c) of the overlapping events exhibit opposite directionality in response to eIF4A3 inhibition. This is consistent with the idea that these AS events are detected as a result of drug-induced eIF4A3 inhibition, which then affects splicing in a manner that is relatively conserved.

**Figure 6:**
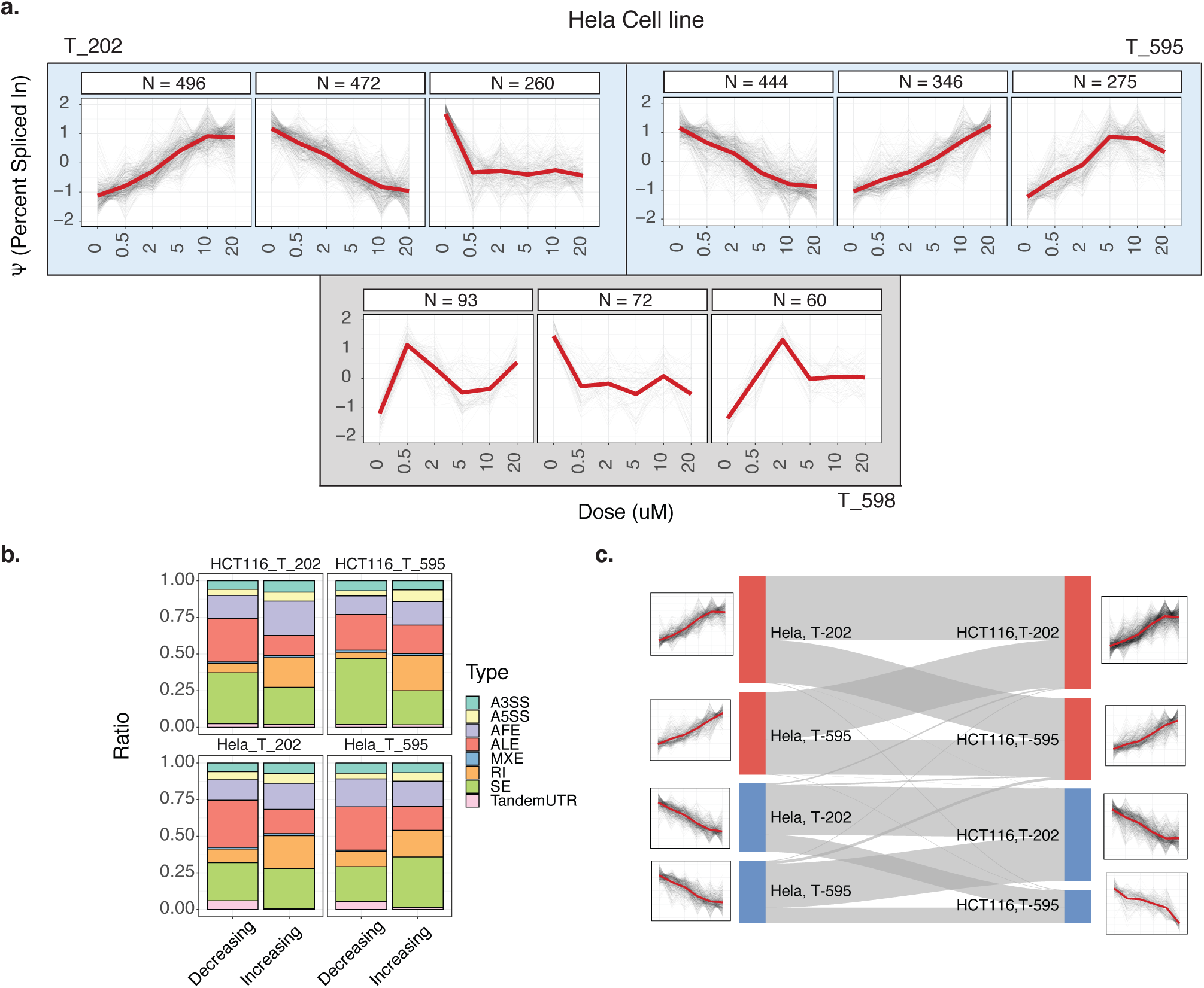
Clustering Ψ response profiles of AS isoforms using WGCNA. **a.** The inclusion levels (Ψ values) of AS isoforms calculated by MISO are clustered using WGCNA. Blue and grey background colors represent the clustering of Ψ profiles for the active compounds and the control compound, respectively (at most 3 clusters are shown per each compound, sorted based on cluster sizes). X and Y axes represent inhibitor concentrations, and MISO Ψ values. Each black line illustrates Ψ response of one MISO isoform in a given condition, and the red lines represent the consensus response in each cluster computed by averaging Ψ values of all the isoforms. **b.** Stacked bars, showing the ratio of each AS type in the set of monotonically increasing and decreasing events. The composition of AS types is clearly different between the two sets. **c.** A sankey diagram illustrating the proportion of shared events between monotonic sets identified in part a. Monotonically increasing and decreasing sets are shown in red and blue rectangles, respectively.

Next, we measured the proportion of AS types in the set of monotonically increasing and the set of monotonically decreasing events, separately. As illustrated in Figure 6b, the composition of AS types is clearly different when the set of events with monotonically increasing Ψ profiles are compared to the events with monotonically decreasing Ψ profiles, while similar proportions are observed in a same set (e.g. monotonically increasing) in the two active inhibitors and the two cell lines. The clearest distinction is observed in RI type, where the proportion is on average 0.135 larger in the set of increasing profiles.

One hypothesis arising from the above is that many of RI affected transcripts would normally undergo NMD, however due to inhibition of eIF4A3 mediated NMD, they escape degradation. To test this idea, we classified AS events into those that cause frameshift based on the local annotations (*i.e*. the skipped exon or retained intron of length not a multiple of 3) and then checked whether monotonic events (union set of MISO and VAST-TOOLS, Supplementary Figure 10) were biased (*Fisher’s exact test*) to be out of frame, compared with background events in MISO or VAST-TOOLS (Materials and Methods section). We observed (Supplementary Table 4) that the set of skipped exons with monotonic responses (both increasing and decreasing) and the set of retained introns only with monotonically increasing responses were significantly enriched by frameshift causing AS events (*p-values* <0.1). Only 15% of both MISO and VAST-TOOLS detected AS events overlapped with genes recognized to be NMD prone in our libraries (Supplementary Figure 6), potentially denoting NMD independent functions of eIF4A3 in AS regulation. Taken together, the data are consistent with the notion that RI (and SE) events are a function of modulation of the two eIF4A3-dependent processes - modulation of AS as well as a block of NMD.

Among AS events, alternative last exons (ALE) also exhibit a clear variation between the set of monotonically increasing and decreasing events (average ratio of 0.165 and 0.289, respectively). Between the two last exons annotated for each ALE event, we denote the one in the 5’ position the “proximal” last exon and the other one the “distal” last exon. For the ALE events shown in Figure 6b, we examined whether the proximal or the distal exon is over-expressed in each set. There is a small (Supplementary Figure 11) yet statistically significant bias towards the proximal exons in the cluster of monotonically decreasing events upon eIF4A3 inhibition by both eutomers. The skewing towards the proximal terminal exons of the genes after eIF4A3 inhibition suggests a role for this gene in terminal exon transcript structure.

Finally, we searched for biological processes whose genes were enriched in the set of genes with AS events in low or high drug concentrations (Supplementary Figure 12, Supplementary Data 18, 19, 20). Biological processes detected in low drug concentrations were almost a subset of biological processes significantly affected in high concentrations of drugs. Differentially spliced genes are involved in similar biological processes in HeLa and HCT116 cell lines including RNA splicing, DNA repair, cell cycle regulation and 3’ end processing. When we did an intersect between the AS pathways, NMD pathways and gene expression pathways, we notice that 81% and 78% of the pathways enriched by AS genes are also targeted through gene expression regulation and NMD regulation (Supplementary Figure 13). On the gene level, we observed little overlap between the genes which exhibit monotonic relationship with eIF4A3 drug doses that are involved in NMD and those that are alternatively spliced (as determined either by MISO or VAST-TOOLS). These findings suggest that the vast majority of eIF4A3 generated AS (particularly SE and RI) does not necessarily generate NMD-prone isoforms as would be expected, although a small subset of genes clearly exhibit this. This is consistent with the idea that eIF4A3 (and hence the EJC) may have independent roles in specific classes of AS events, namely SE, RI and ALE, in addition to its well-known role in NMD.

### Enriched RBP regulatory motifs in 5’ intron regions of eIF4A3 dependent SE and intron regions of eIF4A3 dependent RI events

Finally we set out to determine common features of skipped exons and retained introns inducing AS regulation through eIF4A3 inhibition. To examine the widest range of AS undergoing similar regulation (monotonically increasing *or* monotonically decreasing Ψ profiles), we looked at the union of such events in all 4 drug:cell line pairs of data. For each set of events, all annotated AS events in MISO database not in the set were used as the control set. Considering first structural features, we compared intron and exon lengths of the set of monotonically increasing, monotonically decreasing, and background AS events (Supplementary Figure 14, 15). Exons of the identified monotonically increasing RI events and the identified monotonically decreasing SE events are significantly longer that the background exons (Mann-Whitney U test; p-value <0.01). Also, introns of both groups of monotonic responses are significantly shorter than background events (p-value <0.01).

We next asked whether conserved regulatory features are encoded in eIF4A3 dependent AS sequences, adopting previously described motif enrichment approaches (Ray *et al*, 2013; Funnell *et al*, 2017b). RNA motif density analysis of the most abundant classes, SE and RI AS events, revealed the enrichment of RNA binding protein (RBP) regulatory motifs in 5’ intronic regions of skipped exons for events with both increasing and decreasing Ψ profiles, and intronic regions of retained introns for events with monotonically decreasing responses (Figure 7a, b, Supplementary Data 21, 22, 23, 24). For RNA binding proteins with known binding motifs, the Position Weight Matrices (PWMs) from CIS-BP RNA database (Ray *et al*, 2013) were used. RBPs from the same proteins were grouped together and the enrichment or depletion of them were assessed using Mann-Whitney U test. Motif hits of each set was compared to motif hits of the background set, as explained in more details in Materials and Methods section.

**Figure 7:**
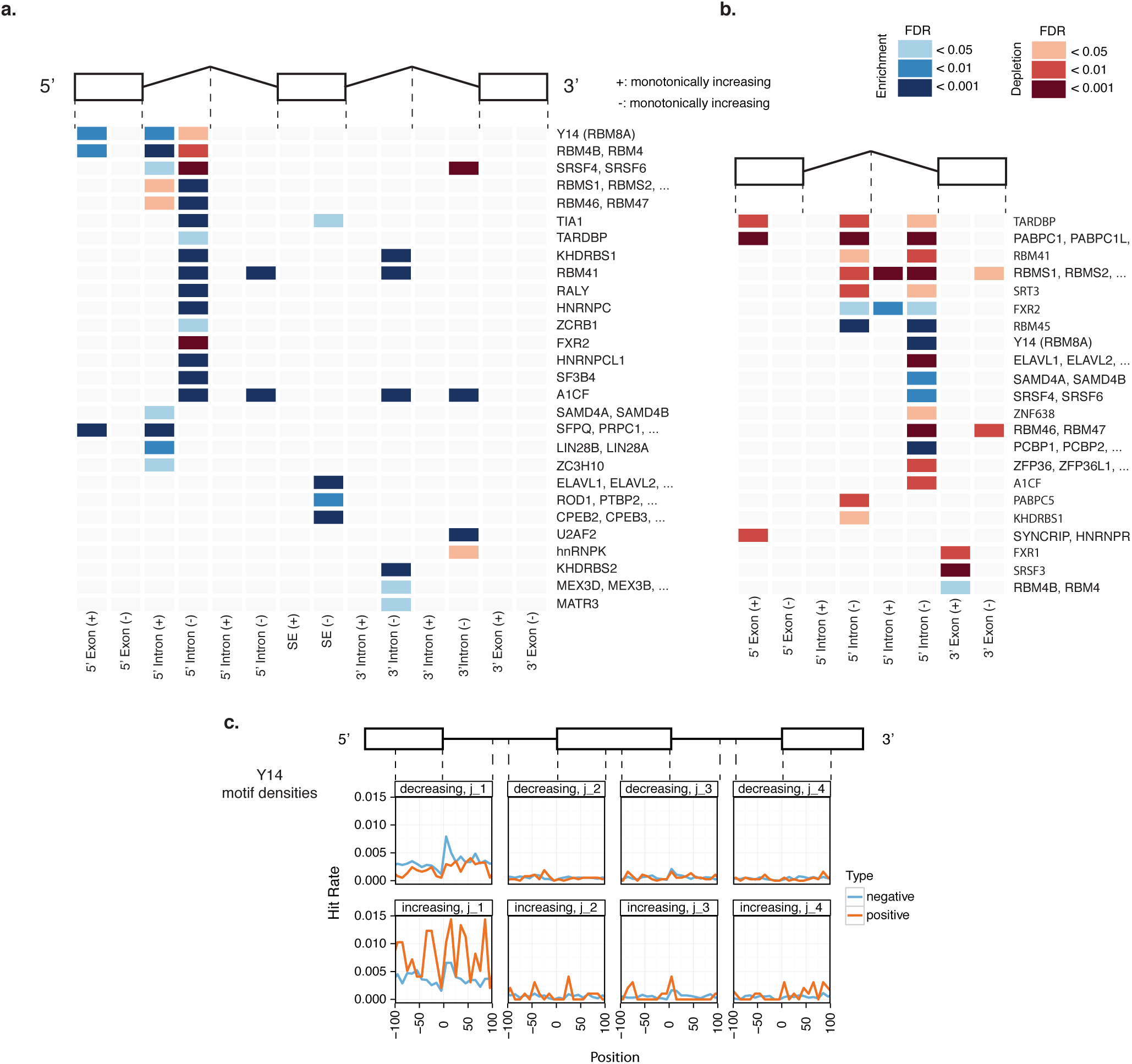
Motifs associated to AS regulation by eIF4A3. **a., b.** Analysis of RNA motifs with known RNA binding proteins. Each row represents a set of RBPs binding to a similar RNA motif, enriched in at least one of the investigated regions around identified AS events. For SE (a.) and RI types (b.) of splicing, exonic regions, and intronic regions adjacent to exon junctions (of length 300) are taken into account. The set of increasing (+) and decreasing (–) monotonic events are separately compared to background AS events. Enriched and depleted motifs are shown in blue and red, respectively. **c.** Motif hits density of Y14, a known member of exon junction complex is shown for 200 nucleotides regions adjacent to SE junctions. For the set of events with increasing and decreasing response profiles normalized hits are compared against negative (background) samples. A clear variation is observed among hit counts of increasing, decreasing, and control sets in the 5’ most junction region.

For SE regions (Figure 7a) both over and under representation of RBP motifs were observed in the 5’ intronic region of monotonically responding gene transcripts, consistent with the known function and location of the eIF4A3 in EJC at and upstream of splice junctions (or exon/intron boundaries). Interestingly, for retained introns (Figure 7b) over or under representation of motifs was predominantly within introns exhibiting monotonically decreasing response profiles although the number of events with decreasing responses was much smaller.

Many of enriched motifs are known to be splicing related, including RBM4, SFPQ, MBNL1-3 (Lai *et al*, 2003; Patton *et al*, 1993; Kalsotra *et al*, 2008). A notable example is Y14/RBM8A (Figure 7c), a core component of 10EJC which functionally cooperates with eIF4A3. This is one of the few RBPs with enriched binding motifs in both 5’ exons and 5’ introns of alternatively spliced exons of increasing responses. Consistently, 5’ intronic regions of events with monotonically decreasing responses are under represented with Y14 binding sites. The distribution of motifs around the splice junction shows a clear variation between these two classes (Figure 7c) in close vicinity of the splice junction, in agreement with Y14’s known functional roles. Taken together, the analysis of monotonic eIF4A3 dependent AS events demonstrates that they may be characterised by longer exons, shorter introns and region specific over/under representation of specific RBP motifs.

## Discussion

Despite the central importance of splicing in multiple disease areas, including cancer, developmental disorders and neuroscience, relatively few small molecule modulators of core spliceosome functions have been described. We have recently described elsewhere (Ito *et al*, 2017b; Iwatani-Yoshihara *et al*, 2017) the synthesis and target specificity (over related helicases) of a series of novel eIF4A3 allosteric inhibitors. Here we show for the first time using these small molecule chemical probes the extent of eIF4A3 dependent transcription and alternative splicing in human cells. To address this we have defined transcripts and alternative splicing reactions that respond in monotonic fashion to increasing short exposure inhibition of eIF4A3, a systems approach to defining the eIF4A3 functional network (Funnell *et al*, 2017a). In addition to revealing dose monotonic transcriptional responses, the availability of a chemically identical but inactive stereoisomer (distomer) of the two active compounds (eutomers) used provides an important control. Very few dose dependent transcript or AS events were observed with the inactive isomer in comparison with active compounds. In addition, we conducted the experiments in two different cell lines, HCT-116 as well as HeLa to control for cell-type specific effects.

Eutomers used in this study allosterically inhibit helicase activity of eIF4A3 (Ito *et al*, 2017a) leading to a reduction in NMD activity measured by reporter plasmids (Iwatani-Yoshihara *et al*, 2017). Treatment of two different cell lines (to control for cell-type specific events) using graded-inhibition has allowed the systematic global identification of genes which are eIF4A3 "dose" dependent (Supplementary Figure 1). When we considered only NMD-prone transcripts in Ensembl, we found that a subset of these NMD-prone transcripts displayed a monotonically increasing relationship with the eutomers when both their expression and specific isoform (PSI) expression were considered (Figure 2). This is consistent with the known involvement of the ATPase activity of eIF4A3 in the EJC formation and in NMD (Nielsen *et al*, 2009; Gehring *et al*, 2009; Wang *et al*, 2014; Iwatani-Yoshihara *et al*, 2017). Most NMD-prone transcripts identified were up-regulated by eutomer treatment reflecting a stabilization of these transcripts by eIF4A3 (and EJC) inhibition; very few transcripts were downregulated as NMD transcripts were normally degraded without drug treatment (Mendell *et al*, 2004). This suggests that eIF4A3-dependent NMD may only affect a subset of NMD-annotated transcripts (Wang *et al*, 2014) and the common set of genes corroborate this (Supplementary Figure 16).

When we surveyed the global cellular processes affected by transcript levels and NMD-prone isoforms when treated with the eutomeric inhibitors of eIF4A3 (Figure 3), we observed NMD-prone transcripts of the genes involved in cell cycle processes such as spindle formation, chromosomal alignment and segregation (DCTN2, NUMA1, CENPV, KIF20A, SKA3, SMC1A, ZWINT), G2/M transition (CCNB1IP1, CDC27, CDK10, SEPT2, PBK), and induction of apoptosis (CCNL2) were affected. It is also significant that a number of genes were involved in the chromosomal passenger complex (CPC) including AURKB and BIRC5 (survivin) as well as downstream components of its signalling pathway such as KIF20A and NSUN2. This implicates eIF4A3 in the maintenance of proper cell cycle regulation particularly through the processes of spindle formation, chromosomal alignment and segregation. This hypothesis was clearly demonstrated in additional experiments where eIF4A3 inhibition, either with compounds or siRNA, revealed G2/M arrest, in conjunction with increased apoptosis, with demonstrable centrosome-spindle disturbances such as mis-segregated chromosomes (Figure 4). This is consistent with previous experiments using high content imaging and siRNA inhibition of eIF4A3 and CDC27 (whose NMD-prone transcript is monotonically stabilised when eIF4A3 is inhibited - Supplementary Figure 17) respectively, resulting in mitotic defects in HeLa cells (Kittler *et al*, 2004). This highlights the notion of evolutionary conservation of these processes (though not the mechanism of EJC involvement in NMD (Gatfield *et al*, 2003)), from Drosophila (Hansen *et al*, 2009) to human.

To ascertain whether the NMD transcripts are normally-spliced and then stabilized (Lewis *et al*, 2003) or whether eIF4A3-dependent alternative splicing may also be involved in generating these NMD-prone transcripts, we examined AS using two complementary tools (MISO and VAST-TOOLS whose annotations databases partially overlap, see Supplementary Figure 5). VAST-TOOLS can identify more retained intron (RI) events, as there are >10 fold more in its splicing database. For skipped exon (SE) events, MISO and VAST-TOOLS share some common events but there are more events specific to VAST-TOOLS than MISO. Alternative First/Last Exon (AFE/ALE) events are predominately identified by MISO. Hence to get the broadest possible range of events, a union set was used (Materials and Methods). The analysis highlights the marked differences in the ALE, RI and SE (only in HCT116) events between the monotonically increasing and decreasing clusters in the two cell lines when treated with both eutomers (Figure 6b), suggesting that eIF4A3 may be involved in these specific AS events. What we observed was the enrichment of SE and RI events in monotonically increasing clusters formed by eutomer treatment that could be predicted to generate pre-mature stop codons (PTCs) by generation of transcripts that are not multiples of three and hence cause frameshifts, assuming the same start frame (Supplementary Table 4).

While the data does not rule out the possibility of NMD isoforms complicating the alternative splicing analysis, it clearly demonstrated that while some AS events may produce NMD transcripts (which are stabilized by inhibition of eIF4A3 and hence the EJC function), all AS events do not necessarily lead to NMD transcripts which are stabilized by eIF4A3 (and EJC) inhibition (Metze *et al*, 2013). This may suggest distinct roles for eIF4A3 and the EJC in the process of splicing and NMD.

Investigation of hard coded sequence features of eIF4A3 dependent AS events demonstrated that introns and exons of transcripts that undergo eIF4A3-dependent AS, particularly RI events and SE events, are longer than the average background (non eIF4A3 dependent) exons and introns (Supplementary Figures 14, 15). We also observed significant variation between ALE in the set of monotonically increasing and decreasing events (Supplementary Figure 11). This finding suggests the involvement of eIF4A3 in the selection of alternative terminal exons of a subset of genes and inhibition of eIF4A3 shows a skewing towards the proximal terminal exons of the genes in the monotonically decreasing clusters, reminiscent of CDK12 action on terminal exons (Tien *et al*, 2017).

A recurrent theme of RNA splicing is the modulation of trans-acting splice accessory factors, RBPs and their cis-acting binding motifs. Having identified the genes that exhibit monotonic relationships to eIF4A3 inhibition, we sought to identify the trans-acting factors by studying the changing patterns of binding motifs of known RBPs in the monotonic RI and SE events (Figure 7). From the analysis it is clear that the motifs of RBPs (e.g. RBM8A/Y14, RBM4B, SRSF4,6, RBM1,2, RBM46,47) depleted in the 5’ intronic region of monotonically increasing SE transcripts are inversely correlated in monotonically decreasing SE transcripts. This mirroring of RBP motifs in two opposing effects are consistent with known mechanisms of splicing and have been previously observed for CLK2 (Funnell *et al*, 2017a). However, unlike CLK2 in which motif pattern changes are observed in both upstream and downstream introns and exons (Funnell *et al*, 2017a), eIF4A3 inhibition results is motif pattern changes predominately in the 5’ intronic regions. Consistent with this, Y14/RBM8A, a core component of EJC which functionally cooperates with eIF4A3 has enriched binding motifs in both 5’ exons and 5’ introns of monotonically increasing transcripts (Figure 7), in close vicinity of the splice junction (Saulière *et al*, 2012), in agreement with Y14’s known functional roles (Nielsen *et al*, 2009).

Finally, our global analysis suggested that eIF4A3 and likely EJC are involved in the NMD of the transcripts of RBPs that have both their NMD transcript and NMD isoform monotonically increase on dose-dependent eutomer treatment (e.g. SRSF2 (Supplementary Figure 18),3,6 (Supplementary Figure 19), RBMX, PSPC1, RBM4B). Many of these RBPs which show differences in binding patterns in the monotonically increasing (e.g. SRSF2,3,6, PPRC1, RBMX, PTBP2) or decreasing (e.g. IGF2BP3, TARDBP, ZFP36, CPEB4, SRSF4, MBNL3) transcripts are themselves modulated by eIF4A3. They illustrate the complexity of the regulatory network and is consistent with the theme of factor auto-regulation in splicing. However, it also underscores the utility in using graded inhibition by eutomers with a distomer control and studying only those genes whose expression displays monotonic relationships to identify targets of eIF4A3.

In conclusion, the use of pharmacologically graded inhibition of eIF4A3 have allowed us to identify a global set of genes involved in chromosome segregation and spindle formation likely responsible for the G2/M mitotic arrest phenotype observed with the active forms of the inhibitor. We further demonstrate that a subset of these genes are NMD-prone and another subset exhibit alternative splicing which point to potentially distinct roles of eIF4A3 and the EJC in NMD and alternative splicing. Finally, we infer from the different motif enrichment and depletion patterns, the RBP factors that might be involved in the EJC-mediated AS and NMD pathways.

## Materials and Methods

### Transfection of siRNAs

Silencer Select siRNAs targeting eIF4A3 were purchased from Life Technologies (s18878: Sense: GCAUCUUG-GUGAAACGUGAtt, Antisense: UCACGUUUCACCAAGAUGCgg; s18876: Sense: GGAUAUUCAGGUUCGU-GAAtt, Antisense: UUCACGAACCUGAAUAUCCaa; s18877: Sense: CGAGCAAUCAAGCAGAUCAtt, Anti-sense: UGAUCUGCUUGAUUGCUCGtt). Twenty four hours after HeLa cells seeding, siRNAs were transfected using Dharmafect 1 (GE Healthcare, Chicago, IL) according to manufacturer’s instructions at a final concentration of 10 nM. Non silencing siRNA was used as the negative control. After 72 h of incubation, cells were harvested for RNA-seq and total RNAs were extracted using an RNeasy Miniprep Kit (Qiagen, Valencia, CA).

### Treatment with compounds

Twenty four hours after HeLa cell seeding, cells were treated with each concentration of compound for 6 h. Cells were harvested for RNA-seq and total RNAs were extracted using an RNeasy Miniprep Kit (Qiagen).

### Apoptosis assay

After 24 h of incubation with the various compounds, caspase-3/7 activity was determined using the Caspase-Glo 3/7 Assay (Promega, Madison, WI) according to the manufacturer’s instructions.

### Immunofluorescence

Cells were fixed in 4% of paraformaldehyde for 15 min at room temperature followed by permeabilization with 0.1% triton buffer. After blocking with 5% BSA, cells were incubated with primary antibody over night at 4 °C. Anti-CENP-B antibody (sc-22788, Santa Cruz, Dallas, TX) and anti-alpha tubulin (T9026, Sigma-Aldrich, St. Louis, MO) were used at concentrations of 0.5 *µ*g/mL and 7.5 *μ*M, respectively. After washing with PBS three times, the cells were incubated with Alexa Fluor 488 anti-mouse IgG or Alexa Fluor 594 anti-rabbit IgG (Life technologies, Carlsbad, CA) at a concentration of 2 *µ*g/mL for 1 h at 37 °C, and were mounted with Vectashield (Vector Laboratories, Burlingame, CA). Cells were imaged on LSM700 (Carl Zeiss, Oberkochen, Germany).

### Alignment and calculating gene and isoform expressions

The RNA-seq libraries in this study consist of 100 nucleotides paired-end unstranded reads from a HiSeq2000. By using the STAR aligner (Dobin *et al*, 2013), the short reads were aligned to the human reference genome (hg19) downloaded from the UCSC genome browser (Meyer *et al*, 2013). Following the alignment step, duplicate reads were removed by employing SAMtools (Li *et al*, 2009; Li, 2011), and the remaining reads were served as the input to the expression and splicing analyses.

Transcript abundances were estimated by using Cufflinks (Trapnell *et al*, 2012). The Cufflinks FPKM (fragments per kilobase of transcript per million reads mapped) values were calculated separately for each library with the annotation file (.gtf) downloaded from Ensembl (GRCh37) (Cunningham *et al*, 2015). The Cufflinks parameters *frag-bias-correct* and *multi-read-correct* were enabled in this step. The expression values of libraries generated by treating the same cell line by different concentrations of the same compound were normalized using the upper quartile normalization method of the edgeR package (Robinson *et al*, 2010; Robinson and Oshlack, 2010), prior to clustering gene and isoform response profiles.

### Alternative splicing analysis

We first used MISO (Mixture of Isoforms) (Katz *et al*, 2010) framework to find AS events when samples treated with compounds were compared to untreated (control) samples. MISO classifies AS events into 8 AS types and for each candidate event, it assigns a Ψ (Percent Spliced In) value (between 0 and 1) representing the inclusion ratio of an isoform in a library. When comparing two conditions, a Bayes Factor (BF) value is reported as a measure of confidence of an event being differentially spliced between the two conditions.

When clustering MISO Ψ values, all reported events were considered. To find AS events induced by compound treatments, treated samples were paired with control samples, and events with |ΔΨ| < 0.1 or BF < 10 were filtered.

In addition to MISO analysis, VAST-TOOLS (Irimia *et al*, 2014; Braunschweig *et al*, 2014) framework was also applied to further explore AS regulation in RNA-seq libraries. VAST-TOOLS classifies AS events into several types and similar to MISO, it calculates Ψ values for events and assesses the difference in these values in two conditions. VAST-TOOLS reports a value indicating the 95% confidence value for the |ΔΨ|, as a measure of significance (MV[abs(dPsi)]: The Minimum Value for |ΔΨ| at 0.95). Events having this value < 0.1 were removed in our analysis.

### Clustering of expression responses

To cluster genes exhibiting correlated response patterns, we applied the WGCNA (Weighted correlation network analysis) framework (Langfelder and Horvath, 2008). The analysis was performed separately for gene expression values, NMD isoform expression values (transcripts annotated as candidates of NMD based on Ensembl annotations), NMD transcripts inclusion levels, and MISO/VAST-TOOLS Ψ values in each set of libraries treated with the same compound and the same cell line. Genes and isoforms with mean FPKM value of < 1 or median value of 0 were removed. Similarly, only genes and isoforms for which the maximum FPKM value was at least 1.5 times larger than the minimum value across compound concentrations were considered; we required some minimum change in abundance values of a gene or an isoform to filter variations due to noise and random RNA-seq sampling. Inclusion levels of NMD isoforms were measured by dividing the isoform expression (FPKM value) by the corresponding gene expression, and was reported as Ψ values. Only isoforms with mean Ψ > 0.05 and median Ψ > 0.001 were kept for clustering. When clustering MISO (VAST-TOOLS) Ψ values, Ψ profiles of all event types were clustered together and no AS events were removed, apart from those filtered by MISO (VAST-TOOLS) itself. In the clustering step, the *networkType* parameter of *blockwiseModules* function was set to “signed”, the *power* parameter was set to 10 for FPKM values and 6 for Ψ values, and for all the other parameters, default values were used.

### Gene set enrichment analysis

Gene set enrichment analysis (GSEA) was carried out on a set of curated GO (Gene Ontology) terms of Biological Processes (BP) from MSigDB database (Liberzon *et al*, 2011). The set of BPs version v.5.2 was downloaded which contains ~4700 GO sets. We applied Hypergeometric test using R (Team, 2014), and *p-values* were corrected using BH (Benjamini and Hochberg) correction for multiple testing.

The EnrichmentMap plugin (Merico *et al*, 2010b) of Cytoscape (Shannon *et al*, 2003; Smoot *et al*, 2011) was used to cluster similar BPs and provide a visual summary of the results. A *FDR* (false discovery rate) cutoff value of 0.01 and a p-value cutoff of 0.005 were applied to the identified enriched GO sets, and Jaccard Coefficient was used as a measure of similarity of GO sets with a cutoff value of 0.25. Names assigned to clusters were manually curated based on the set of BPs in clusters.

### Motif analysis

We searched for known motifs of RNA-binding proteins (RBPs) in different regions of genes that undergo AS regulation through inhibiting eIF4A3, based on the identified RI (retained intron) and SE (skipped exon) events with monotonic Ψ response profiles of MISO and VAST-TOOLS analysis. MISO and VAST-TOOLS data bases were merged and repetitive events were removed.

For RI and SE events constituting two isoforms, the isoforms were grouped into several regions. For SE events we consider 7 regions: 5’ exon, skipped exon, 3’ exon, 300 nucleotides from the start/end of the 5’ intron, and 300 nucleotides from the start/end of the 3’ intron. Similarly, 4 regions are investigated for RI events: 5’ exon, 3’ exon, and 300 nucleotides from the start/end of the retained intron.

Positive samples in our analysis comprise all events with monotonic response profiles in MISO or VAST-TOOLS in any of the 4 active compound:cell line data sets. For the motif enrichment analysis, these evens were divided into two groups: monotonically increasing and monotonically decreasing ones, as the mechanism of regulation could be different.

The set of all events with monotonically increasing (decreasing) Ψ profiles were compared to the set of background MISO/VAST-TOOLS events not identified in any of the 4 active compound:cell line data sets to determine the enrichment or depletion of RNA motifs with known RNA-binding proteins. Positive and negative samples were split into bins of similar lengths and for each positive sample, 10 negative samples from the same bin were selected.

Position Weight Matrices (PWMs) for binding motifs of RBPs were downloaded from CIS-BP RNA data base (Ray *et al*, 2013). First, the background frequencies of A/C/G/U nucleotides were determined. Next, for a given motif and a candidate sequence, the log odds score of that sequence being generated randomly based on background nucleotide frequencies was compared to it being generated based on PWM weights. Then, the maximum value for these log odds scores for each motif was computed and all the sequences with scores above 80% of the maximum score were counted as hits for the corresponding motif.

All motifs sharing a same RBP were clustered together and their hits were merged. Finally, hit counts were normalized by regions’ lengths, Mann-Whitney U test was applied to compare hit counts in regions of positive events compared to hit counts in background events; *p-values* were calculated and corrected using BH multiple test correction method. Only motifs for which the frequency of hits in positive regions were at least 25% higher or at least 25% lower than background regions were considered.

### Frameshift analysis

We investigated the potential of the identified RI and SE events to induce frameshifts. For the VAST-TOOLS and MISO clustering results of active compounds, events with monotonic Ψ response profiles were grouped together (2 groups, increasing and decreasing). Considering that two isoforms are reported in MISO/VAST-TOOLS annotation files for each event, if the length of a retained intron or an skipped exon is not a multiple of 3, the event can induce a frameshift based on the local evidence. For the union of events with increasing (decreasing) response profiles in MISO and VAST-TOOLS, a control set was formed by all the events in MISO/VAST-TOOLS knowledge bases other than those in the set, and the ratios of events that can cause frameshifts were compared between the two sets using *Fisher’s exact test*.

### Data Availability

Raw sequence reads used in this study are available at the Short Read Archive (SRA) under the identifier SRP117312 and the BioProject identifier PRJNA401938. All other data available from the authors upon reasonable request.

